# A High-throughput Multi-Species Platform Using Biolayer Interferometry Immunosorbent Assay (BLI-ISA) as an Alternative to Indirect ELISA for Vaccine Development

**DOI:** 10.1101/2025.07.24.666607

**Authors:** Ti Lu, Md Shafiullah Parvej, Sean K. Whittier, Suhrid Maiti, Zackary K. Dietz, Debaki R. Howlader, Mst Nusrat Zahan, Satabdi Biswas, William D. Picking, Wendy L. Picking

**Author notes:** Co-corresponding Authors: Wendy L Picking,; Ti Lu,.

## Abstract

In vaccine development, the ELISA (Enzyme-Linked Immunosorbent Assay) is commonly used to compare the antibody titers of samples from several treatment groups. This often requires extensive sample preparation, manual labor, and long incubation and processing times. Biolayer Interferometry (BLI) has emerged as an alternative to the ELISA for the detection and quantification of antigen-specific antibodies in biological samples. However, the implementation of BLI as a replacement for the ELISA in vaccine development requires that experimental parameters are established for accurate and reproducible results. Here we give a general protocol for a biolayer interferometry immunosorbent assay (BLI-ISA) for the comparison of antigen-specific antibody levels in treatment group sera that uses secondary antibody binding responses as replacement for ELISA endpoint titers. We also validate that this BLI-ISA yields the same results as the ELISA endpoint titer while requiring far less time and effort.

## Introduction

Traditional methods for analyzing vaccine efficacy include various forms of the enzyme-linked immunosorbent assay (ELISA, see **Supplemental Table S1**), a well-established plate-based assay used in immunology for detecting and quantifying antibodies (Sakamoto, Putalun et al. 2018). Indirect ELISA provides a robust platform for detecting specific antibodies through interaction with an immobilized antigen followed by adding a secondary antibody-enzyme conjugate that allows for an amplified colorimetric readout (Lequin 2005). Frequently, antigen-specific antibodies in sample sera are quantified in terms of arbitrary “ELISA Units”, which report how much a given sample must be diluted to give a signal at some level above the limit of detection (Lequin 2005, Wang, Bashiri et al. 2024).

While effective, the method is labor-intensive and time-consuming (Lequin 2005). Prior to performing the ELISA, antigen must be adsorbed to wells in ELISA plates, after which the plates must be blocked to limit non-specific antibody binding, a process which takes several hours using a limited number of samples (Hosseini, Vázquez-Villegas et al. 2018). For endpoint titer determination, serum samples must be serially diluted, which can require a significant time and labor investment when many samples are to be compared (Frey, Di Canzio et al. 1998). Finally, performing the assay requires aliquoting reagents and samples into the plate wells, followed by numerous washing and incubation steps, before the plates are read in a plate reader. The entire procedure can easily take up a researcher’s entire day and each pipetting step can introduce errors (Hosseini, Vázquez-Villegas et al. 2018).

Biolayer Interferometry (BLI), on the other hand, is an optical biosensing technology that can replace the ELISA by measuring changes in the wavelength of light reflected from the biosensor surface caused by molecular binding in real time. This eliminates the need for enzymatic detection schemes and enables rapid antibody detection (Petersen 2017). BLI eliminates the need to coat and wash plates because the antigen is specifically and rapidly bound to the biosensor surface, and the detection of primary and secondary antibody binding is monitored in real-time over the course of several minutes (Sun, Reid et al. 2013, Markwalter, Jang et al. 2017).

This Biolayer Interferometry Immunosorbent Assay (BLI-ISA) can provide the same information as the ELISA in a fraction of the time (Markwalter, Jang et al. 2017, Dzimianski, Lorig-Roach et al. 2020). However, prior to its employment as an ELISA replacement, experimental parameters of the BLI-ISA must be determined such that there is a linear correlation between the BLI-ISA result and ELISA endpoint titer for any given sample. As we have discovered, this process of experimental tuning can take a significant amount of time and may be a deterrent to some groups hoping to replace the ELISA. Here we provide a general BLI-ISA protocol for the determination of antigen-specific antibody levels in vaccine treatment groups that reproduces ELISA results in a fraction of the time. Furthermore, this BLI-ISA uses the BLI binding response value as a proxy for ELISA endpoint titers rather than binding rates or initial slopes, eliminating the need for curve fitting in data analysis.

### General BLI-ISA outline

The BLI-ISA as described here uses Ni-NTA biosensors to specifically bind His-tagged antigen. After antigen binding, biosensors are briefly washed in buffer before being transferred to wells containing sample serum that has been diluted in binding buffer. Primary antibodies are allowed to bind for a predetermined time, followed by a wash step. Biosensors are then transferred to wells containing a secondary antibody conjugated with horseradish peroxidase (HRP). Secondary antibodies are allowed to bind for a predetermined time and antibody levels are quantified as the optical wavelength shift at the end of this period (**Figure 1, Table 1**). It must be noted that secondary antibodies do not have to be HRP-conjugated and there is no need for an HRP substrate. An unmodified secondary antibody can also be used, however, the HRP adds additional mass that increases the resulting BLI signal.

**Table 1.** Side-by-side comparison of the time duration.

| Step | BLI-ISA | Indirect ELISA |
| --- | --- | --- |
| Sensor/Plate Preparation | Ni-NTA biosensor hydration (10 min). | Plate coating with antigen (overnight or 2–4 hrs at 37°C). |
|  |  | Blocking with milk (1–2 hrs). |
| Baseline and Antigen Loading | Baseline stabilization (4 min). | Antigen immobilization is completed during plate coating. |
|  | Antigen loading (8–10 min). |  |
| Sample Incubation | Serum detection (8–10 min). | Serum incubation (1–2 hrs at 37°C). |
|  |  | Washing steps between incubations. |
| Secondary Antibody Detection | Secondary antibody binding (6–7 min). | Incubation with HRP-conjugated secondary antibody (1–2 hrs). |
|  | Washing steps are automated. | Multiple washing steps (manual). |
| Signal Detection | Real-time binding response detected immediately during the experiment. | TMB substrate addition (10–15 min), reaction stop, and absorbance reading. |
| Total Duration | 30–40 minutes per set of serum (3 samples and one reference with duplicated for individual). | 4–6 hours (including overnight coating for some protocols). |

**Figure 1.**
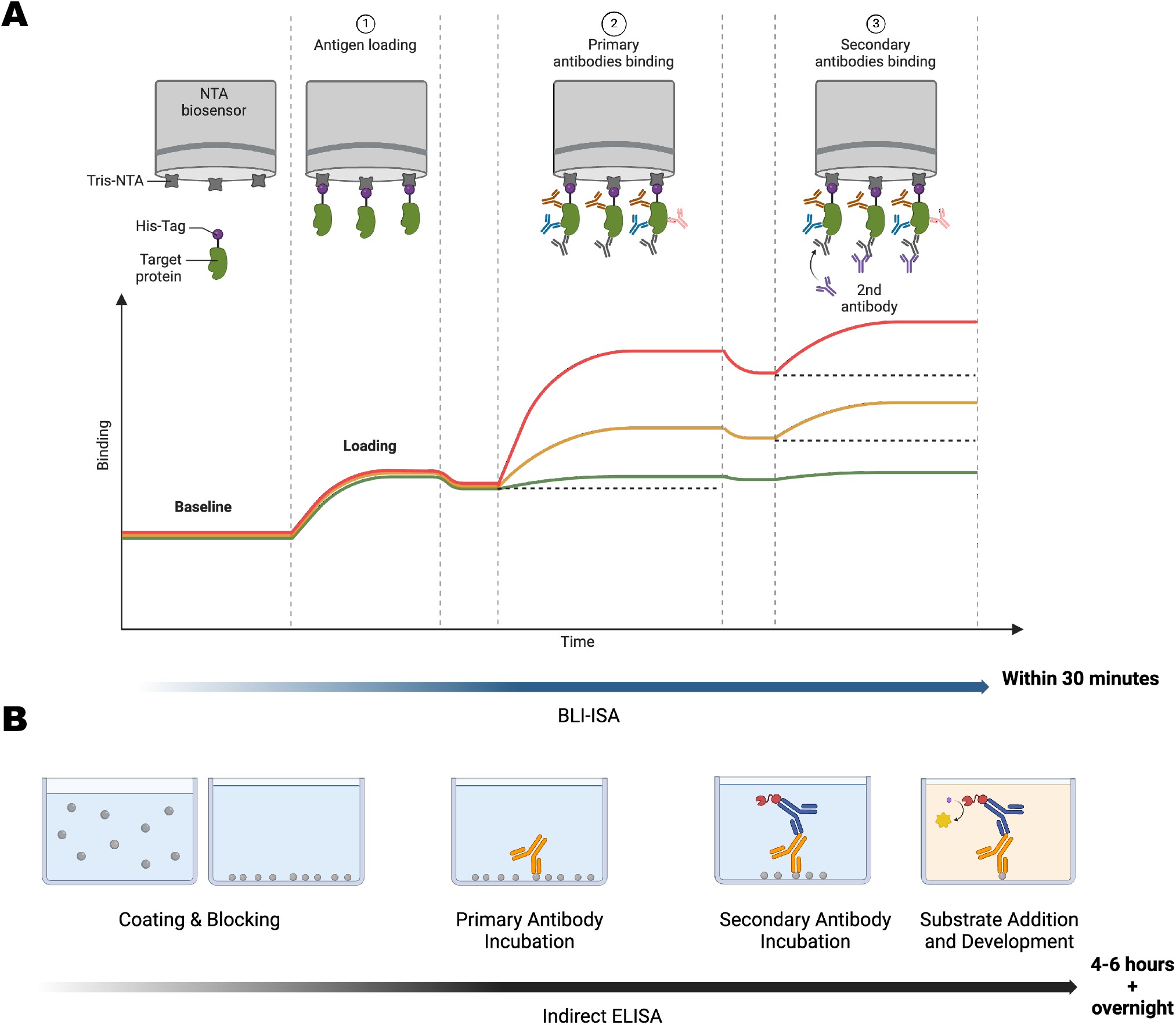
Workflow and representative binding curves of BLI-ISA. The diagram illustrates the steps of Biolayer Interferometry Immunosorbent Assay (BLI-ISA) and corresponding binding curves over time. **Top:** Schematic representations of each step highlight the molecular interactions occurring on the sensor surface. **Middle:** Binding response curves for different serum samples are shown during the antigen loading, primary antibody binding, and secondary antibody binding steps. **Bottom:** A comparison of the analogous steps and durations required for Indirect ELISA, including antigen coating, blocking, primary antibody incubation, and secondary antibody detection using enzyme-conjugated antibodies.

### Required Materials

To perform the BLI-ISA (Biolayer Interferometry - Immunosorbent Assay), a suitable Octet instrument (ideally with 8 sensors), compatible biosensors, and specified 96-well plates will be needed. Essential reagents include recombinant His-tagged antigens and serum samples, both of which must be diluted or prepared in assay buffer optimized for Ni-NTA capture of His-tagged antigens (**Table 2**).

**Table 2.** The materials and instruments of the BLI-ISA assay.

| Category | Item | Details |
| --- | --- | --- |
| Instrumentation | Suitable Octet BLI Instrument | Ideally an Octet instrument with at least 8 sensors for efficient data collection. |
|  | Biosensors | Ni-NTA biosensors for binding of His-tagged antigen. |
|  | Specified BLI Plates | Black, flat-bottom 96 well plates (Greiner part # 655209). |
| Reagents | His-Tagged Antigen | Antigen should be prepared in assay buffer (composition provided below) at a concentration of 20 µg/ml. |
|  | Serum Samples | Serum to be tested, diluted with assay buffer sufficiently to fall within linear range of BLI binding response (see Results and Discussion). |
|  | Secondary antibody | HRP-conjugated secondary antibody against serum host species primary antibodies, diluted to 12.5 µg/ml in assay buffer. |
|  | Assay Buffer | A suitable buffer such as TBS (Tris-buffered saline, pH 8) supplemented with 0.1% Tween-20 to reduce non-specific binding. |

### Plate set-up and biosensor preparation

The 96-well plate for BLI-ISA is organized as follows (**Figure 2**): columns 1, 3, 10, and 12 (grey wells) contain assay buffer for pre-wetting and equilibrating biosensors, column 2 (green wells) holds the diluted His-tagged antigens for loading onto the sensors, columns 4–9 contain serum samples or analytes to be tested (red wells, rows A-F) and serum from negative control groups (purple wells, rows G-H) for reference subtraction, and column 11 (yellow wells) contains the secondary antibody for detection. Biosensors are pre-wet in column 1, loaded with protein in column 2, washed in columns 3, 10, and 12, and systematically interact with serum samples in columns 4–9, followed by secondary antibody binding in column 11. The biosensor tray is carefully loaded into the instrument’s sensor holder, and the 96-well assay plate is aligned in the plate holder. Instrument settings are configured to match the plate layout, ensuring appropriate agitation speeds (1000 rpm) and precise alignment during each step. This setup ensures efficient preconditioning, protein loading, washing, sample interaction, secondary antibody binding, and regeneration for robust assay performance.

**Figure 2.**
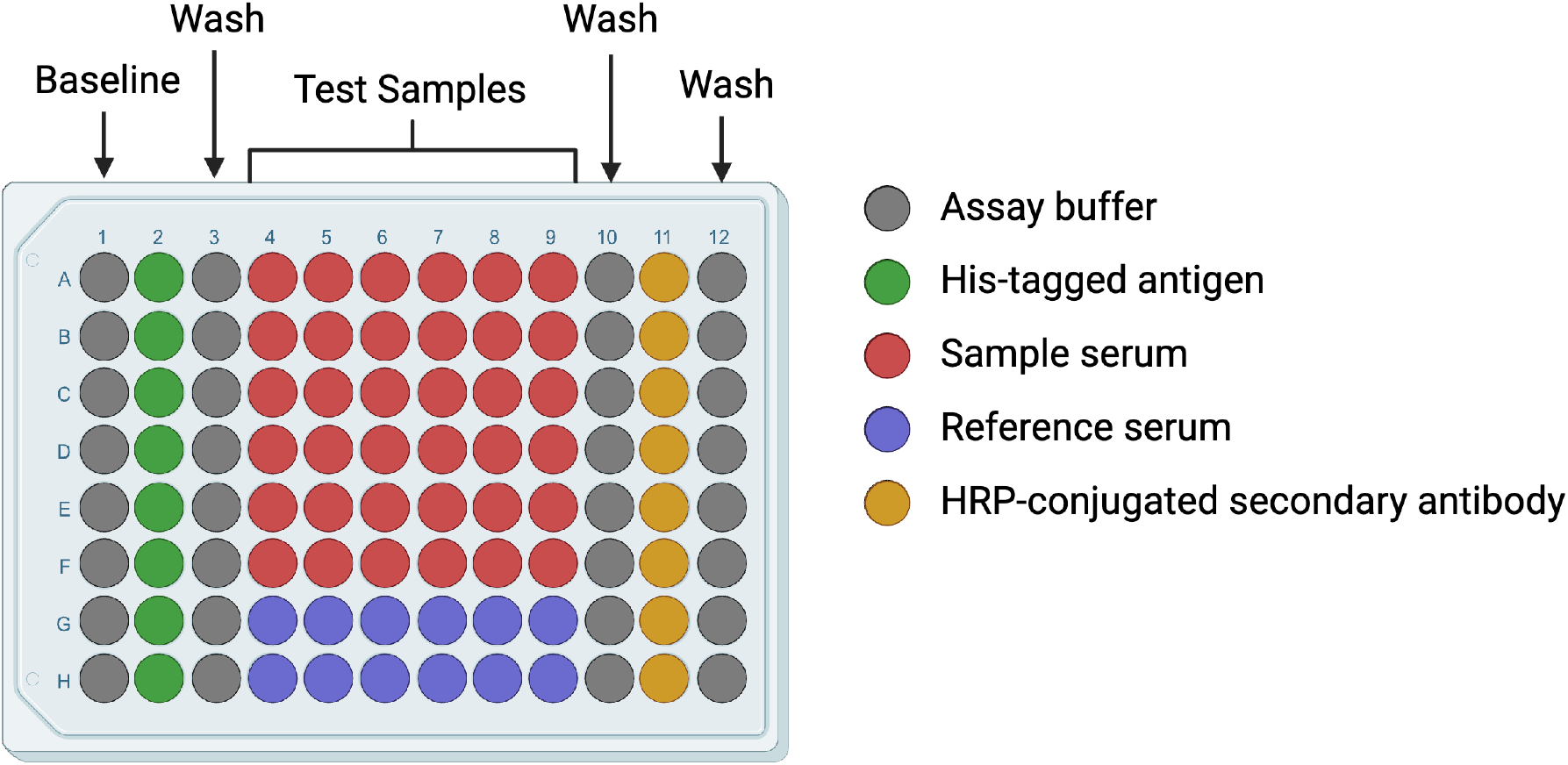
Plate Layout for BLI-ISA Analysis. The 96-well plate layout used for the BLI-ISA experiment is shown, highlighting the categorization of different sample groups. Gray wells represent buffer-only wells for baseline readings and wash, green wells correspond to dilute His-tag antigen proteins, and red wells contain test serum samples for evaluation. Purple wells are designated as negative control sera for background subtraction, while yellow wells represent HRP-conjugated secondary antibodies.

### Assay steps

The assay is conducted at a plate temperature of 30°C and all steps utilize a shaking speed of 1000 rpm. The start of the experiment is delayed for 10 minutes to allow the plate to thermally equilibrate. The experimental procedure then follows a six-step workflow (sample data are shown below for determining linear responses):

1. <u>Baseline Measurement:</u> Sensors are transferred to assay buffer located in Column 1 and incubated for 240 seconds.
2. <u>Protein Loading:</u> The sensors are moved to a solution of His-tagged antigen in assay buffer in Column 2 for 500 seconds to immobilize antigen onto the sensors.
3. <u>Post-Loading Baseline:</u> The sensors are transferred to assay buffer in Column 3 and incubated for 240 seconds.
4. <u>Primary antibody loading:</u> Sensors are transferred to sample serum diluted with assay buffer in one of the sample Columns 4-9 for 500 seconds (refer to **Figure 2**).
5. <u>Post-Serum Baseline:</u> Sensors are moved to assay buffer in Column 10 and incubated for 240 seconds.
6. <u>Association Phase:</u> Sensors moved to anti-IgG secondary antibody diluted with assay buffer in Column 11 and incubated for 400 seconds.

The above procedure is repeated using a fresh set of biosensors for each of the remaining sample Columns 4-9. Each column corresponds to a different serum sample, with every two wells testing a specific dilution of that sample. This approach ensures that measurements fall within the linear range of BLI-ISA response values, which is essential for accurate comparison with ELISA endpoint titers.

### Data processing

Antibody responses in serum samples were analyzed using the Octet BLI Analysis Studio 13.0 software. All sensor data were aligned along the Y-axis at the start of the association step. For reference subtraction, sera from negative control wells (purple wells in the plate layout, **Figure 2**) were designated as reference wells. The averaged reference well signal is then subtracted from each sample well’s association phase data to correct for non-specific binding and dissociation of His-tagged antigen. The antigen-specific IgG response was quantified as the binding response in nanometers (nm) of wavelength shift at the conclusion of the 400-second association step.

## Results

### BLI-ISA gives similar results as ELISA

BLI-ISA demonstrated comparable results to a standard indirect ELISA, with a strong overall correlation between results from each assay (**Figure 3**). Species-specific binding responses were robust across all tested groups (mice, rats, rabbits, and pigs), with BLI-ISA binding results correlating well with ELISA antibody titers (R^2^ values ranging from 0.7326 to 0.9768, **Figure 3A-D**). There is, however, an observed nonlinearity between BLI-ISA results and ELISA endpoint titers for samples with high BLI binding responses. For example, while mouse sample 5 showed a 3-times higher ELISA titer than mouse sample 4, it exhibited only a 1.3-fold greater binding response in BLI-ISA (**Figure 3A** and **Supplemental Table S2**). This indicates that while the methods are directionally consistent, BLI-ISA may have a compressed linear range compared to ELISA.

**Figure 3.**
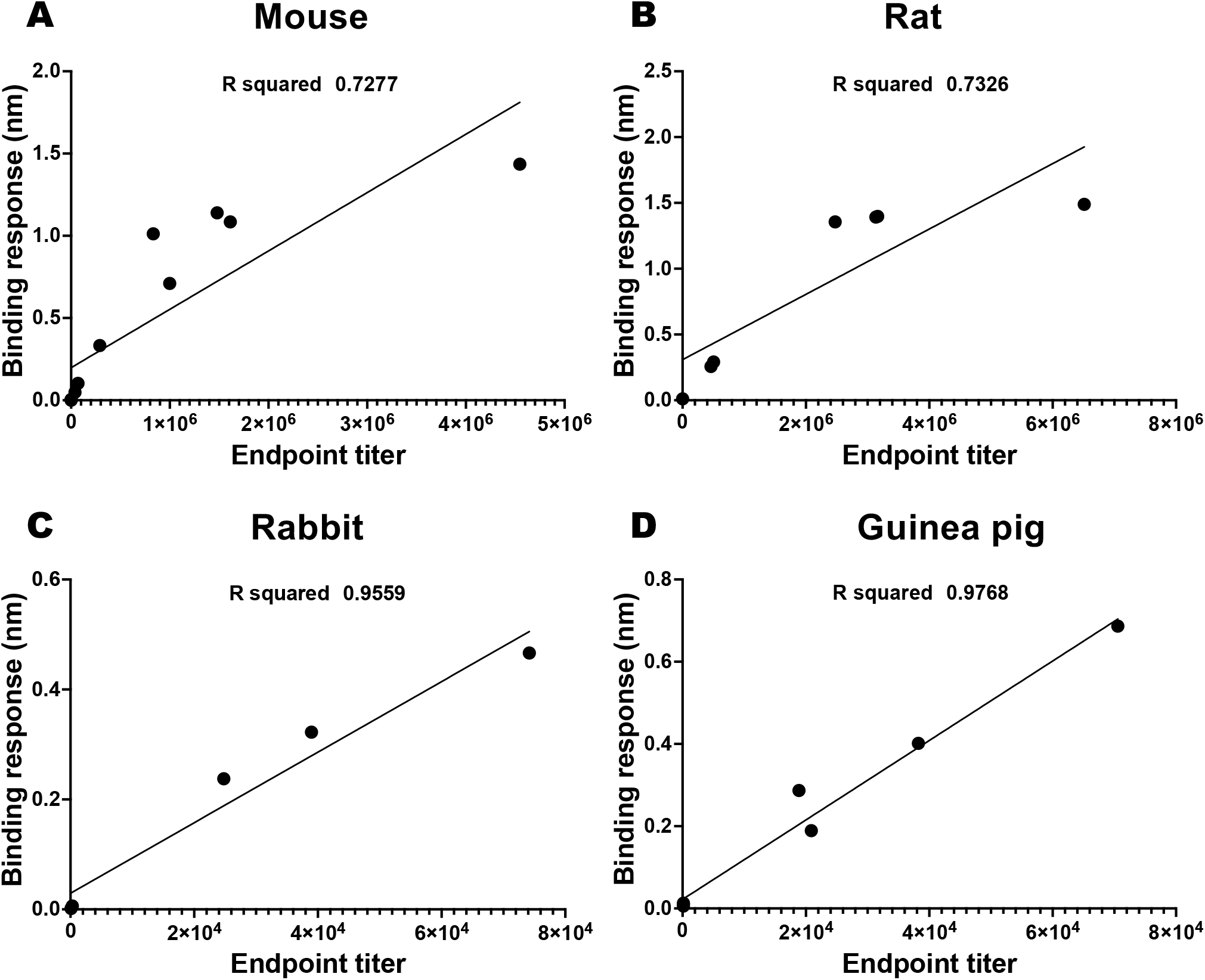
Correlation between BLI-ISA binding responses and ELISA endpoint titers for each specie. (A–D) Linear regression analysis of BLI-ISA binding responses (nm) and ELISA endpoint titers for sera from mice (A), rats (B), rabbits (C), and guinea pigs (D). High correlation coefficients (R^2^) demonstrate the strong agreement between BLI-ISA and ELISA measurements across species. In mice and rats, however, the higher titer sera may require adjustments regarding dilution.

### Linear response range of BLI-ISA

To improve the correlation between BLI-ISA and ELISA results, we determined the linear range of BLI-ISA response values using a dilution series of two different sample sera containing antibodies that contain antibodies against the *Pseudomonas* protein PcrV (**Figure 4**) or antibodies against the *Shigella* protein IpaB (**Supplemental Figure S1**). Binding responses scaled proportionally with antibody concentrations in the range of 0.25 to 1.5 nm, as indicated by strong linear regression in that region (R^2^ = 0.9919, **Figure 4B;** R^2^ = 0.9801, **Supplemental Figure S1**). Limiting our mouse and rat BLI-ISA data to samples that are within this linear range dramatically improves the correlation with ELISA endpoint titers, with R^2^ values increasing from 0.7277 and 0.7326 to 0.9292 and 0.9776, respectively (**Figure 4C** and **D**). It is therefore important that sample sera dilutions are adjusted to stay within this linear response range to ensure comparable results to ELISA.

**Figure 4.**
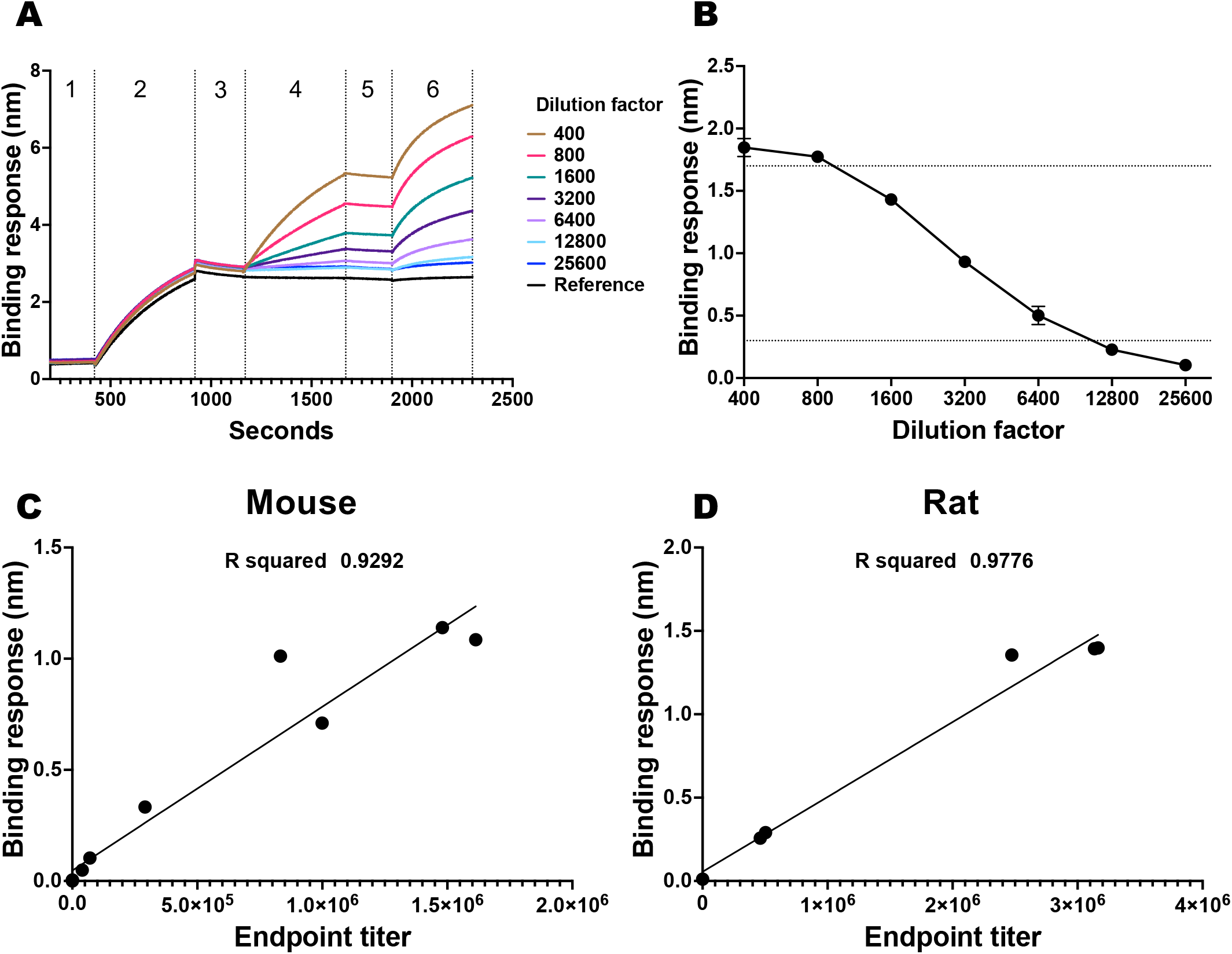
Serial diluted binding curves for BLI-ISA. (A) Real-time binding responses of His-tagged PcrV (*P. aeruginosa* type 3 secretion system antigen) with serially diluted rat-vaccinated serum samples, measured over time. The curves demonstrate consistent binding signals across dilutions ranging from 1:400 to 1:25600. (B) Binding responses at equilibrium plotted against serum dilutions, showing a proportional relationship between binding response and dilution factor. Linear regression analysis of BLI-ISA binding responses (nm) and ELISA endpoint titers for sera from mice (C) and rats (D) after excluding outliners. High correlation coefficients (R^2^) demonstrate the strong agreement between BLI-ISA and ELISA measurements across species.

## Discussion

Our study demonstrates that BLI-ISA can provide comparable results to ELISA while dramatically reducing processing time. For the same number of samples, BLI-ISA required only 30–40 minutes due to its streamlined workflow, real-time detection, and automated washing steps, as opposed to the 4–6 hours required by ELISA, which involves manual washing, extended incubations, and an overnight antigen coating step (**Table 1** and **Figure 1**). Furthermore, BLI-ISA uses observed BLI binding response values to compare specific antibody levels, rather than binding rates or initial slopes. This eliminates the need for curve fitting or linear regression, greatly improving data analysis. Overall, BLI-ISA provides a versatile and scalable platform for antigen-antibody interaction studies. This makes it particularly well-suited for accelerating vaccine development workflows, enabling faster candidate evaluation without compromising accuracy and reproducibility (Petersen 2017).

Critical to the implementation of BLI-ISA is the need for the binding response values to vary linearly with antibody levels. Our results indicate that this is true for response values in the 0.25-1.5 nm range. Experiments utilizing BLI-ISA should plan to dilute serum samples accordingly, so that the final response value is within this range. This may require that some samples be tested at different dilution values. In general, samples with expected low levels of specific antibodies should be diluted in the 200- to 500-fold range, while samples with high antibody titers should be diluted in the 1000- to 2000-fold range. Response values can then be corrected for differential dilution provided they are within the linear range.

In addition to saving time, it is important to evaluate the cost implications of BLI-ISA. While biosensor trays used in the Octet system incur a higher upfront cost compared to ELISA reagents, they significantly reduce the labor time involved in sample processing (Dzimianski, Lorig-Roach et al. 2020). For Ni-NTA sensors, a key limitation is that they may need to be replaced after every set of eight samples due to reduced binding capacity following regeneration, thereby increasing the cost for large-scale experiments. However, this cost is offset by the reduction in labor costs, particularly for groups that perform frequent ELISAs and/or ELISAs for many samples (Markwalter, Jang et al. 2017).

It is worth mentioning that alternative biosensor options are available for capturing antigens, such as SA (streptavidin) for biotinylated antigens or amine-reactive biosensors (AR2G) for covalent linkage of antigen to the biosensor surface (Bhalla, Jolly et al. 2016, Huang, Xie et al. 2022). These sensors permanently bind antigens, eliminating the slow signal drift observed when His-tagged antigen dissociates from Ni-NTA biosensors (Bhalla, Jolly et al. 2016). Furthermore, because SA and AR2G sensors completely immobilize the target antigen, trays of biosensors can be pre-loaded with antigen and stored in compatible buffer for minimally several days prior to conducting the BLI-ISA. This allows the the antigen loading step to be eliminated when testing samples, further reducing the processing time (Castro, Bezerra et al. 2022, Jiang, Han et al. 2024). On the other hand, these approaches require protein modification, such as biotinylation or covalent attachment to the biosensor at exposed primary amines, which can potentially alter the antigen’s tertiary structure or block epitopes critical for antibody recognition (Fowler, Evers et al. 2011). Thus, while these sensors offer advantages in stability and time savings, careful consideration of their impact on the antigen/antibody interaction is required.

## Materials and Methods

### Materials

Chromatography columns were from GE Healthcare (Piscataway, NJ). NTA Biosensors were from Sartorius (Bohemia, NY). All other reagents were from Sigma or Fisher Scientific and were chemical grade or higher.

### Protein production

Production of IpaB and PcrV has been described in detail previously (Das, Howlader et al. 2020, Lu, Howlader et al. 2023). Briefly, plasmids expressing His-IpaB (cloned from *Shigella flexneri*) or His-PcrV (cloned from *Pseudomonas aeruginosa*) were used to transform *E. coli* Tuner (DE3) cells. After growth in a 10L bioreactor, protein expression was induced with 1mM IPTG and the culture was grown at 37°C for three hours. Bacteria were collected, resuspended in IMAC binding buffer, lysed, and the supernatant was subjected to standard IMAC purification using a Ni-NTA column (Martinez-Becerra, Chen et al. 2013, Lu, Das et al. 2023). Fractions were eluted using a gradient of elution buffer (containing up to 500mM imidazole), pooled, and further purified via a Q FF column using a linear salt gradient. The final samples were concentrated, buffer-exchanged to TBS with 0.05% LDAO. All proteins had LPS levels <5 EU/mg as determined using the Endosafe nexgen system (Hu, Varisco et al. 2023).

### Serum Samples

Mouse, rabbit, guinea pig, and rat sera were used to evaluate the binding responses across different species. Pooled serum samples used in this study were obtained from on-going vaccine studies, where all animals were vaccinated as part of separate research projects. In one study, animals were vaccinated using a nanoemulsion formulation containing the *Pseudomonas aeruginosa* protein PcrV (Howlader, Das et al. 2021) and in the other study animals were vaccinated using a nanoemulsion containing the *Shigella flexneri* protein IpaB (Lu, Raju et al. 2024). These studies employed well-characterized vaccination protocols, and all serum samples were collected following standardized immunization and serum harvesting procedures, ensuring consistency across samples (Das, Howlader et al. 2020, Lu, Howlader et al. 2023). Detailed information can be found in the Supplemental **Table 2**. Ethical approval for all animal experiments was obtained from the respective Institutional Animal Care and Use Committees (IACUC) 38241 at the University of Missouri.

### IgG ELISAs

Pooled anti-IpaB or anti-PcrV IgG titers were measured as previously described (Das, Howlader et al. 2020, Lu, Howlader et al. 2023). Briefly, microtiter wells were coated with 100 ng IpaB or PcrV in 100 µl PBS, incubated at 37°C for 3 hours, and blocked with 10% nonfat dry milk in PBS overnight. Sera were added as the primary antibody for 2 hours, followed by an HRP-secondary IgG antibody (1:1000) for 1 hour. After washing, OPD substrate was added, and detection was done at 490 nm via ELISA. Endpoint titers were calculated and represented as ELISA units per ml (EU ml^-1^). All tests repeated twice.

### Data Analysis

Data processing and statistical analyses were conducted using GraphPad Prism version 8.1.2. Simple linear regression was employed to identify the linear response range and to calculate correlation coefficients between BLI-ISA and ELISA data. Analyses were performed with 95% confidence intervals and automatic determination of regression ranges, ensuring robust quantitative comparisons between the two methods.

## Author contributions

T.L. led data analysis, data generation, visualization,, and drafted the initial manuscript; W.L.P. oversaw conceptualization, secured funding, managed the project, and provided overall supervision; M.S.P contributed to data acquisition, curation, and initial analysis; S.K.W. conducted conceptualization, method development, detailed statistical analyses, and critically reviewed and edited the manuscript; S.T., D.R.H, S.B., and M.N.Z. helped to prepare samples; Z.K.D. handled specimen collection, protein purification, ME nanoemulsion preparation, and animal method development; W.D.P. finalized manuscript editing and writing.

## Acknowledgements

This work was funded by NIAID grants R01AI138970 and R01AI169781 to WLP.

## Data Availability Statement

All datasets generated and analyzed during this study are included in the manuscript and supplementary materials.

