## Supplemental Table S1 for "A High-throughput Multi-Species Platform Using Biolayer Interferometry Immunosorbent Assay (BLI-ISA) as an Alternative to Indirect ELISA for Vaccine Development"

**Supplemental Table S1. Acronyms used in this paper.**

| | |
|---|---|
| BLI | Biolayer interferometry |
| ELISA | Enzyme-linked immunosorbent assay |
| BLI-ISA | Biolayer interferometry immunosorbent assay |
| SA | Streptavidin |
| LDAO | Lauryl-dimethylamine oxide |
| MOPS | 3-(N-morpholino)propanesulfonic acid |
| IPTG | Isopropyl β-D-1-thiogalactopyranoside |
| IMAC | Immobilized Metal Affinity Chromatography |
| EU | Endotoxin Unit |

**Supplemental Table S2. Summary of the BLI-ISA binding responses and ELISA titers of each species.**

| Project & Species | BLI | | ELISA titration | Target antigen | Source |
|---|---|---|---|---|---|
| | Binding response | Dilution factor | | | |
| Serum Dilution-Rat | 1.847 - 0.103 | 400-25600 | 6511646.7 | His-PcrV | In this lab |
| Serum dilution-mouse | 1.232 - 0.039 | 1000-64000 | 6494726.92 | His-IpaB | In this lab |
| Mouse-1 | 0.004107 | 1000 | 44.3 | His-PcrV | In this lab |
| Mouse-2 | 0.004719 | 1000 | 60.2 | | |
| Mouse-3 | 0.006422 | 1000 | 45.2 | | |
| Mouse-4 | 0.8749 | 1000 | 1613529 | | |
| Mouse-5* | 1.167 | 1000 | 4546266 | | |
| Mouse-6 | 0.0066 | 500 | 15 | | |
| Mouse-7 | 0.0539 | 500 | 39857.6 | | |
| Mouse-8 | 0.9845 | 500 | 832473.6 | | |
| Mouse-9 | 0.3524 | 500 | 290728.8 | | |
| Mouse-10 | 1.094 | 500 | 1480417.8 | | |
| Mouse-11 | 0.1211 | 500 | 70180.2 | | |
| Mouse-12 | 0.8206 | 500 | 998973 | | |
| Rat-1 | 0.2907 | 1000 | 503417.7 | His-PcrV | In this lab |
| Rat-2 | 0.2569 | 1000 | 461808.6 | | |
| Rat-3 | 1.3926 | 1000 | 3135234.2 | | |
| Rat-4* | 1.4887 | 1000 | 6511646.7 | | |
| Rat-5 | 1.3558 | 1000 | 2474395.6 | | |
| Rat-6 | 1.3978 | 1000 | 3164154.7 | | |
| Rat-7 | 0.0112 | 1000 | 13 | | |
| Rabbit-1 | 0.3223 | 100 | 38964.7 | His-PcrV | In this lab |
| Rabbit-2 | 0.2377 | 100 | 24788.9 | | |
| Rabbit-3 | 0.001669 | 100 | 94.1 | | |
| Rabbit-4 | 0.4664 | 100 | 74186 | | |
| Rabbit-5 | 0.006026 | 100 | 308.2 | | |
| G-Pig-1 | 0.6864 | 200 | 70561.9 | His-IpaB | In this lab |
| G-Pig-2 | 0.2868 | 200 | 18870 | | |
| G-Pig-3 | 0.01288 | 200 | 135.5 | | |
| G-Pig-4 | 0.4015 | 200 | 38234.4 | | |
| G-Pig-5 | 0.1892 | 200 | 20884 | | |
| G-Pig-6 | 0.01033 | 200 | 140.4 | | |
| G-Pig-7 | 0.005687 | 200 | 152.2 | | |

*: Outlier that excluding for $R2$ modification.

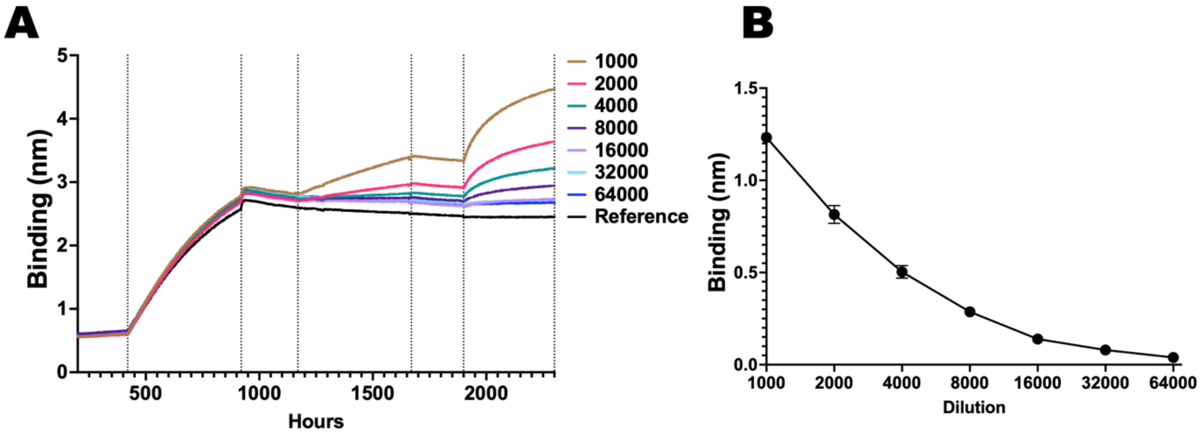


**Supplemental Figure 1.** Serum dilution analysis with His-IpaB (from *S. flexneri*) and mouse-vaccinated serum. (**A**) Representative binding curves are shown for serum dilutions ranging from 1:1000 to 1:64000. They show clear association and saturation phases. (**B**) Linear regression analysis of binding responses in the range of 0.2 to 1.2 nm ($R^2 = 0.9801$) is presented, highlighting the linear relationship between response and serum dilution.
